# Mechanism of microtubule plus-end tracking by the plant-specific SPR1 protein and its development as a versatile plus-end marker

**DOI:** 10.1101/521047

**Authors:** Rachappa Balkunde, Layla Foroughi, Eric Ewan, Ryan Emenecker, Valeria Cavalli, Ram Dixit

**Author notes:** **Corresponding Author** Ram Dixit, Biology Department, Washington University, One Brookings Drive, Campus Box 1137, Saint Louis, MO 63130, USA.

## Abstract

The dynamics and functions of microtubule plus-ends are governed by microtubule plus-end tracking proteins (+TIPs). Here, we report that the diminutive *Arabidopsis thaliana* SPIRAL1 (SPR1) protein, which regulates directional cell expansion, is an autonomous +TIP. Using *in vitro* reconstitution experiments and total internal reflection fluorescence microscopy, we demonstrate that the conserved N-terminal domain of SPR1 and its GGG motif are necessary for +TIP activity, whereas the conserved C-terminal domain and its PGGG motif are not. In addition, we show that the N-and C-terminal domains, either separated or in tandem, are sufficient for +TIP activity and do not significantly perturb microtubule plus-end dynamics compared to full-length SPR1. We also found that exogenously expressed SPR1-GFP and NC-GFP label microtubule plus-ends in animal cells. These data establish SPR1 as a new type of intrinsic +TIP and demonstrate its utility as a universal microtubule plus-end marker.

## INTRODUCTION

Microtubules are dynamic polymers of αβ-tubulin heterodimers which form different arrays to orchestrate critical cellular activities in eukaryotes. The plus-ends of microtubules are of particular significance because they drive the assembly and disassembly of microtubules and connect microtubule tips to various cellular structures. Specialized proteins called microtubule plus-end tracking proteins (+TIPs) specifically accumulate at growing microtubule plus-ends and control their behavior and interactions. +TIPs can either directly or indirectly target growing microtubule plus-ends (Akhmanova and Steinmetz, 2010). For example, the evolutionarily conserved End Binding 1 (EB1) protein autonomously targets growing microtubule plus-ends by recognizing GTP-bound tubulin structures (Bieling et al., 2007; Dixit et al., 2009; Maurer et al., 2011; Maurer et al., 2014; Maurer et al., 2012). In turn, EB1 recruits a host of other proteins as part of a dynamically changing microtubule plus-end complex (Honnappa et al., 2009; Kumar et al., 2017).

Land plants contain a unique set of +TIPs (Nakajima et al., 2004; Sedbrook et al., 2004; Wong and Hashimoto, 2017), perhaps because they assemble morphologically and functionally distinct microtubule arrays compared to animals (Young and Bisgrove, 2011). One of the plant-specific +TIPs is the *Arabidopsis thaliana* SPIRAL1 (SPR1) protein. *SPR1* was identified through a genetic screen for skewed root growth (Furutani et al., 2000) and encodes a 12 kDa protein that localizes to the growing plus-end of cortical microtubules and contributes to anisotropic cell expansion (Nakajima et al., 2004; Nakajima et al., 2006; Sedbrook et al., 2004). SPR1 genetically and physically interacts with the *A. thaliana* End Binding 1b (EB1b) protein (Galva et al., 2014), which tracks growing microtubule plus-ends similar to animal and yeast EB proteins (Dixit et al., 2006; Komaki et al., 2010). However, SPR1 is also able to bind to free tubulin and the microtubule lattice independently of EB1b (Galva et al., 2014), raising the question of whether SPR1 needs EB1b for microtubule plus-end localization. Here, we report that SPR1 is an autonomous +TIP and identify the domains and critical residues that are required for this activity. Finally, we show that a fusion protein consisting of the conserved N-and C-terminal domains of SPR1 fused to GFP labels growing microtubule plus-ends in animal cells, thus making it a universal microtubule plus-end marker.

## RESULTS AND DISCUSSION

### SPR1 is an autonomous microtubule plus-end tracking protein

To study the mechanism of SPR1’s +TIP activity, we took an *in vitro* reconstitution approach with purified recombinant SPR1-GFP protein (Figure 1A) and dynamic microtubules. We found that SPR1-GFP on its own labels microtubule plus-ends exclusively during the growth phase (Figure 1B, 1C; Movie 1). Furthermore, SPR1-GFP expressed under its native promoter labels microtubule plus-ends in *eb1a;eb1b* double mutant (Figure 1D), demonstrating that SPR1 is an autonomous +TIP. Given that SPR1 is structurally distinct from other +TIPs (Akhmanova and Steinmetz, 2010; Young and Bisgrove, 2011), this finding establishes a new category of intrinsic +TIP unique to plants.

**Figure 1.**
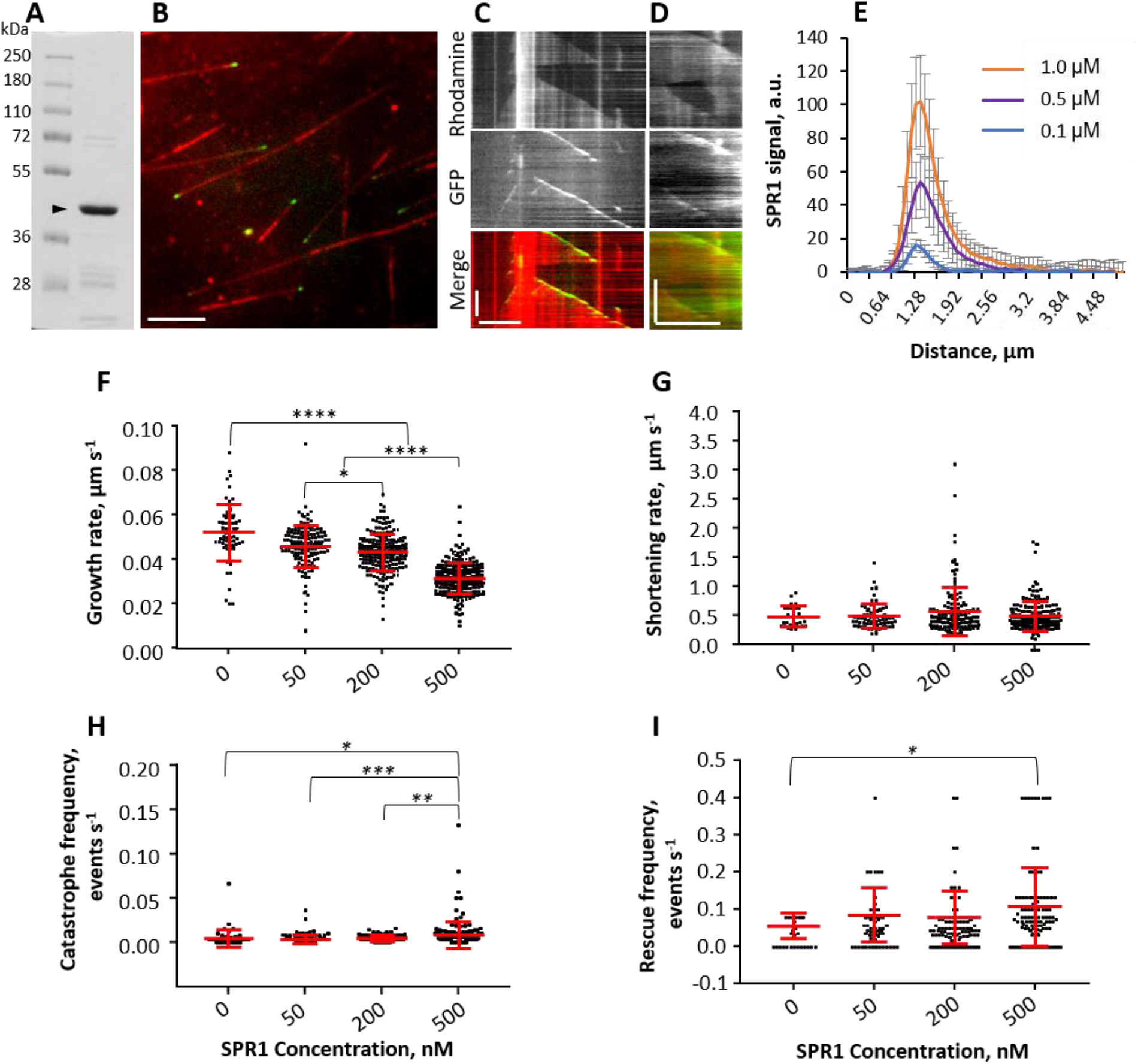
SPR1 is an autonomous microtubule plus-end tracking protein. (A) Coomassie-stained gel of recombinant SPR1-GFP protein. The arrowhead indicates the expected size (40 kDa) of the protein. Numbers on the left indicate size of the protein bands in the ladder. (B) Representative image of SPR1-GFP (green) and dynamic rhodamine-labeled microtubules (red). Scale bar, 5 μm. (C) Representative kymograph showing that SPR1-GFP (green) specifically localizes to the growing plus-end of a microtubule (red). X-axis scale bar, 5 μm; Y-axis scale bar, 100 s. (D) Representative image of SPR1-GFP (green) and RFP-TUB6 (red) in *eb1a;eb1b* double mutant. X-axis scale bar, 5 μm; Y-axis scale bar, 100 s. (E) Graph of SPR1-GFP signal intensity at the plus-end of growing microtubules. Values represent mean ± S.D. n = 52, 89 and 133 for 0.1 μM, 0.5 μM, and 1 μM of SPR1-GFP, respectively. (F-I) Dot plots of microtubule growth rate (F), shortening rate (G), catastrophe frequency (H) and rescue frequency (I) at the indicated SPR1-GFP concentrations. Statistical analyses were conducted using one-way ANOVA. Asterisks indicate significant difference: ****p< 0.0001, ***p< 0.001, **p< 0.01 and *p< 0.05)

We observed that SPR1-GFP also weakly labels growing microtubule minus-ends *in vitro* (Figure 1C; Movie 1), similar to yeast and mammalian EB1 proteins (Bieling et al., 2007; Dixit et al., 2009). The significance of the minus-end tracking activity of SPR1 is not clear. One possibility is that SPR1 regulates the localization of plant microtubule minus-end proteins similar to the regulation of microtubule minus-end binding of CAMSAP2 by human EB1 and EB3 (Yang et al., 2017). In plants, the SPR2 protein localizes to and stabilizes microtubule minus-ends (Fan et al., 2018; Leong et al., 2018; Nakamura et al., 2018). The *spr1-1;spr2-1* double mutant greatly exacerbates the microtubule-dependent growth defects of either single mutant (Furutani et al., 2000), but it remains to be seen whether SPR1 affects SPR2’s localization.

The SPR1-GFP signal is concentrated at the tips of growing microtubules and decays exponentially along the microtubule shaft in a comet-like pattern (Figure 1B, 1E). Increasing the concentration of SPR1-GFP protein enhanced the accumulation of SPR1 at growing microtubule plus-ends (Figure 1E). These results indicate that SPR1 has higher affinity for growing microtubule plus-ends compared to the microtubule lattice.

To determine whether SPR1-GFP affects the behavior of microtubule plus-ends *in vitro*, we measured the dynamic instability parameters at various SPR1-GFP concentrations. We observed that SPR1-GFP inhibited the growth rate of microtubule plus-ends and enhanced their catastrophe frequency in a concentration-dependent manner (Figure 1F, 1H). In contrast, SPR1-GFP did not significantly affect the shortening rate of microtubule plus-ends and only modestly increased their rescue frequency at the highest SPR1-GFP concentration (Figure 1G, 1I). These observations (summarized in Supplemental Table 1) are largely consistent with the reported changes in microtubule plus-end dynamics in *spr1* loss-of-function mutants compared to wild-type (Galva et al., 2014; Lindeboom et al., 2018).

### The conserved N-terminal domain is the primary determinant of SPR1’s +TIP activity

SPR1 is the smallest known +TIP and we wanted to identify the domains that confer microtubule plus-end localization. The conserved N- and C-terminal domains of SPR1 (Figure 2A) have been proposed to contribute to microtubule binding because they contain a GGG and PGGG motif, respectively, which are part of the microtubule-binding repeats of the MAP2/tau family of microtubule-associated proteins (Nakajima et al., 2004; Sedbrook et al., 2004). Individually, the N- and C-terminal domains of SPR1 do not rescue *spr1* mutants and do not bind to microtubules *in vivo* (Nakajima et al., 2004). However, a fusion protein consisting of GFP flanked by the N- and C-terminal domains of SPR1 (N-GFP-C) was partially functional and localized to microtubule plus-ends *in vivo* (Nakajima et al., 2004), suggesting that both domains are needed to recognize growing microtubule plus-ends.

**Figure 2.**
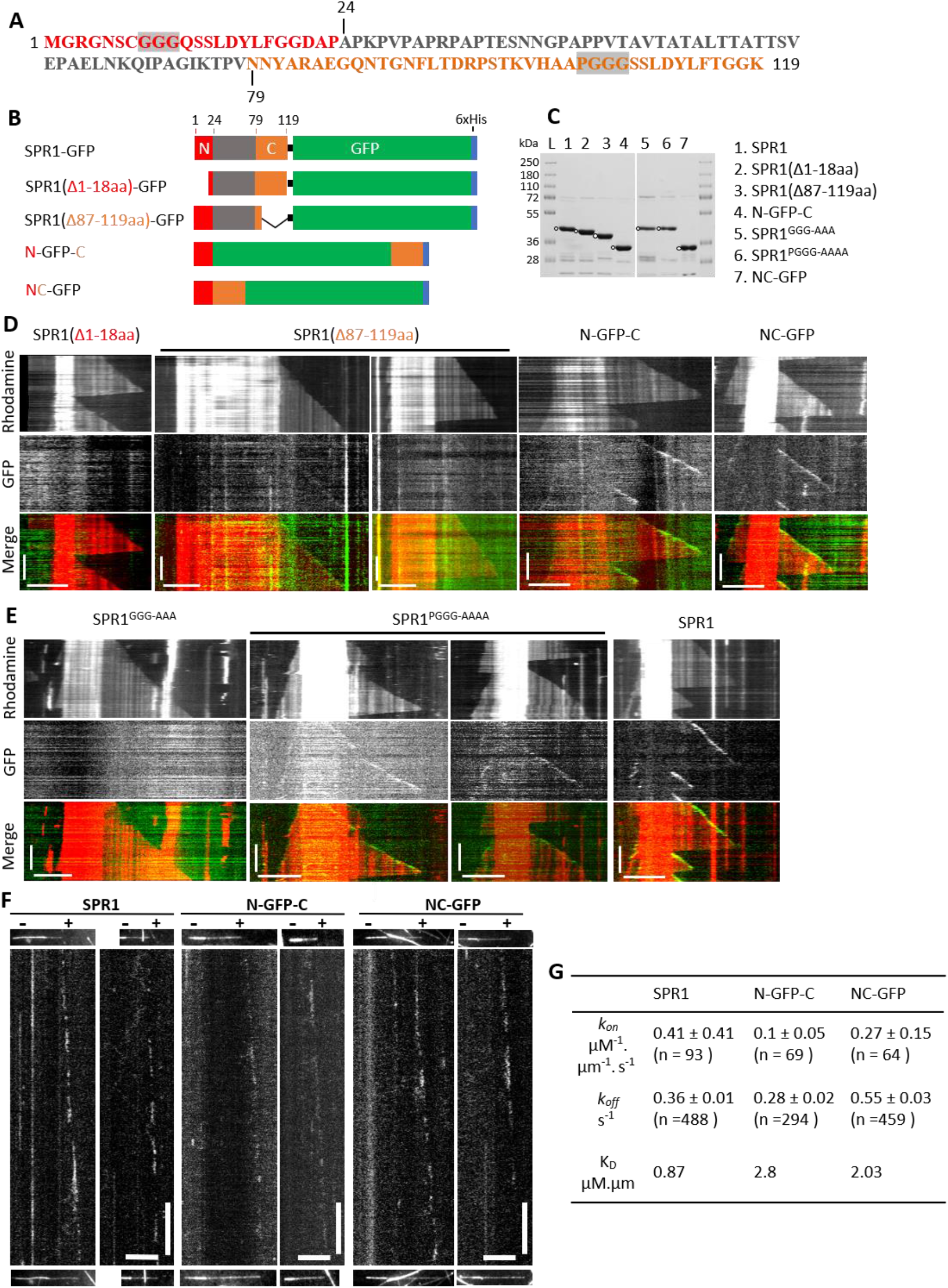
N-terminal domain of SPR1 is required for microtubule plus-end tracking. (A) SPR1 amino-acid sequence. The conserved N-terminal and C-terminal domains are shown in red and orange, respectively. The central variable region is in grey. Grey boxes indicate the conserved GGG and PGGG motifs in the N-terminal and C-terminal domains, respectively. (B) Schematic presentation of the full-length SPR1-GFP fusion protein and mutant SPR1 proteins used in this study. Color code is the same as in (A). (C) Coomassie-stained gel of the truncated and chimeric versions of SPR1 protein used in this study. White dots indicate the expected protein bands. (D) Representative kymographs of SPR1(Δ1-18)-GFP, SPR1(Δ87-119)-GFP, N-GFP-C and NC-GFP proteins. X-axis scale bar, 5 μm; Y-axis scale bar, 100 s. (E) Representative kymographs of SPR1^GGG-AAA^-GFP, SPR1^PGGG-AAAA^-GFP proteins. A kymograph of wild-type SPR1-GFP is shown for comparison. X-axis scale bar, 5 μm; Y-axis scale bar, 100 s. (F) Representative kymographs from single molecule imaging experiments of SPR1-GFP, N-GFP-C and NC-GFP proteins. Images at the top and bottom of the kymographs show the microtubule at the beginning and at the end of the streaming-mode image acquisition. X-axis scale bar, 5 μm; Y-axis scale bar, 10 s (G) Table of the binding rate constant (*k_on_*), unbinding rate constant (*k_off_*) and apparent dissociation constant (*K_D_*) of SPR1-GFP, N-GFP-C and NC-GFP proteins. Values are mean ± S.D. (n = number of molecules)

To determine the relative contribution of the N-and C-terminal domains of SPR1, we created truncated versions of SPR1 that lacked either the N-terminal or the C-terminal domain (Figure 2B, 2C). We found that deleting the N-terminal 18 amino acids (SPR1Δ1-18) abolished +TIP activity *in vitro* (Figure 2D), indicating that the N-terminal region of SPR1 is essential to recognize the microtubule plus-end. Deletion of the C-terminal 33 amino acids (SPR1Δ87-119) diminished the signal at growing microtubule plus-ends but did not eliminate it (Figure 2D), indicating that the C-terminal region enhances the +TIP activity of SPR1 but is not necessary for it. Consistent with these findings and with previous *in vivo* work (Nakajima et al., 2004), the chimeric N-GFP-C protein clearly localized to growing microtubule plus-ends *in vitro* (Figure 2D, Movie 2). To test whether the spacing of the N-and C-terminal domains is important for +TIP activity, we fused these domains in tandem to GFP (Figure 2B and 2C). The resulting NC-GFP fusion protein also localized to growing microtubule plus-ends *in vitro* (Figure 2D, Movie 3).

During these experiments, we noticed that the microtubule plus-end signal of N-GFP-C and NC-GFP were considerably less than the SPR1-GFP signal. To determine whether the N-GFP-C and NC-GFP proteins have a lower affinity for the microtubule plus-end compared to SPR1-GFP, we conducted *in vitro* dynamic microtubule assays under single-molecule imaging conditions followed by kymograph analysis (Figure 2F). Our measurements revealed differences in the binding rate constant *(k_on_)* and unbinding rate constant *(koff)* between the three proteins for the growing microtubule plus-end which yielded a 2-3 fold lower apparent dissociation constant (*K_D_*) for N-GFP-C and NC-GFP compared to SPR1-GFP (Figure 2G). Whether these differences are due to the spacing between the N- and C-terminal domains being different within N-GFP-C and NC-GFP compared to SPR1-GFP or because the variable internal region of SPR1 somehow contributes to microtubule binding remains to be determined.

Next, we tested the significance of the N-terminal GGG and C-terminal PGGG motifs by mutating these amino acids to alanine within the context of full-length SPR1 (Figure 2B, 2C). While mutation of PGGG to AAAA (SPR1^PGGG-AAAA^) had no effect on the +TIP activity of SPR1, mutation of GGG to AAA (SPR1^GGG-AAA^) abolished this activity (Figure 2E). These findings underscore the essential role of the SPR1 N-terminal domain and indicate that the GGG motif is critical for +TIP activity.

### The N-GFP-C and NC-GFP markers do not significantly alter microtubule dynamics

The autonomous +TIP activity of N-GFP-C and NC-GFP make them attractive microtubule plus-end markers. To determine whether they affect microtubule plus-end dynamics similar to the wild-type SPR1 protein, we tested the effect of N-GFP-C and NC-GFP on microtubule dynamics *in vitro.* We found that N-GFP-C did not significantly alter the microtubule growth rate (Figure 3A), catastrophe frequency (Figure 3C) and rescue frequency (Figure 3D), and only modestly increased the shortening rate (Figure 3B). NC-GFP did not significantly alter any of these microtubule dynamic instability parameters (Figure 3E-3H and Supplemental Table 1). Furthermore, we compared the number of transition events exhibited by individual microtubules at different concentrations of SPR1-GFP, N-GFP-C and NC-GFP. We found that SPR1-GFP boosted microtubule transitions in a concentration-dependent manner (Figure 3I). In contrast, neither N-GFP-C nor NC-GFP significantly altered the number of microtubule transitions (Figure 3J and 3K). Taken together, our findings demonstrate that N-GFP-C and NC-GFP localize to growing microtubule plus-ends without significantly perturbing their dynamics unlike wild-type SPR1. These data also provide a potential explanation for why N-GFP-C only partially compliments the *spr1-6* mutant even though it localizes to microtubule plus-ends *in vivo* (Nakajima et al., 2004).

**Figure 3.**
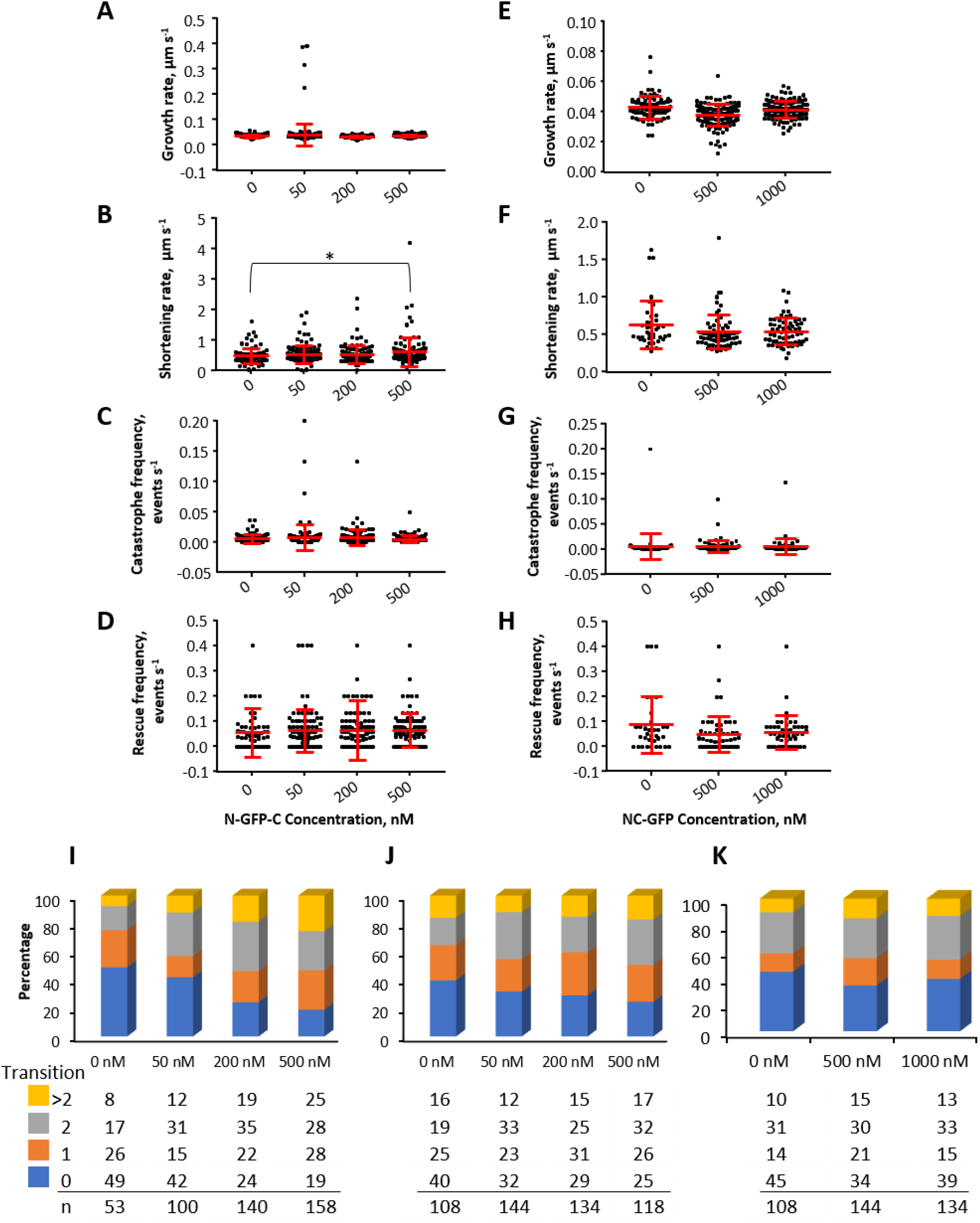
N-GFP-C and NC-GFP do not significantly affect microtubule dynamics *in vitro.* (A-D) Dot plots of microtubule growth rate (A), shortening rate (B), catastrophe frequency (C) and rescue frequency (D) at the indicated N-GFP-C protein concentrations. Asterisks indicate significant difference between the indicated data sets determined by one-way ANOVA, *p< 0.05. (E-H) Dot plots of microtubule growth rate (E), shortening rate (F), catastrophe frequency (G) and rescue frequency (H) at the indicated NC-GFP protein concentrations. (I-K) Stacked bar graphs of the percentage of microtubule transition events at increasing concentrations of SPR1-GFP (I), N-GFP-C (J), and NC-GFP (K). Numbers below show the percentages of each category of microtubule transitions during the observation period.

### SPR1-GFP and NC-GFP label growing microtubule plus-ends in animal cells

Since SPR1-GFP, N-GFP-C and NC-GFP labeled the plus-ends of microtubules assembled from porcine tubulin *in vitro*, we reasoned that they might label microtubule plus-ends in animal cells. To test the suitability of these proteins as microtubule plus-end markers in animal cells, we introduced them into Human Embryonic Kidney-293 (HEK-293) cells and imaged the transfected cells using total internal reflection fluorescence microscopy. Interestingly, both SPR1-GFP and NC-GFP clearly labeled growing microtubule plus-ends and the microtubule shaft to a lesser extent (Figure 4A, 4B; Movie S4 and Movie S5). Although N-GFP-C showed extensive cytoplasmic signal in HEK-293 cells, we were unable to detect unambiguous microtubule plus-end labeling Microtubule growth rate measured by tracking SPR1-GFP and NC-GFP comets showed a growth rate of 0.39 ± 0.14 μm s^-1^ (n = 35) and 0.18 ± 0.04 μm s^-1^ (n = 27), respectively. These values are comparable to microtubule growth rates reported in various cell lines (Goulimari et al., 2008; Sehrawat et al., 2011; van der Vaart et al., 2011).

**Figure 4.**
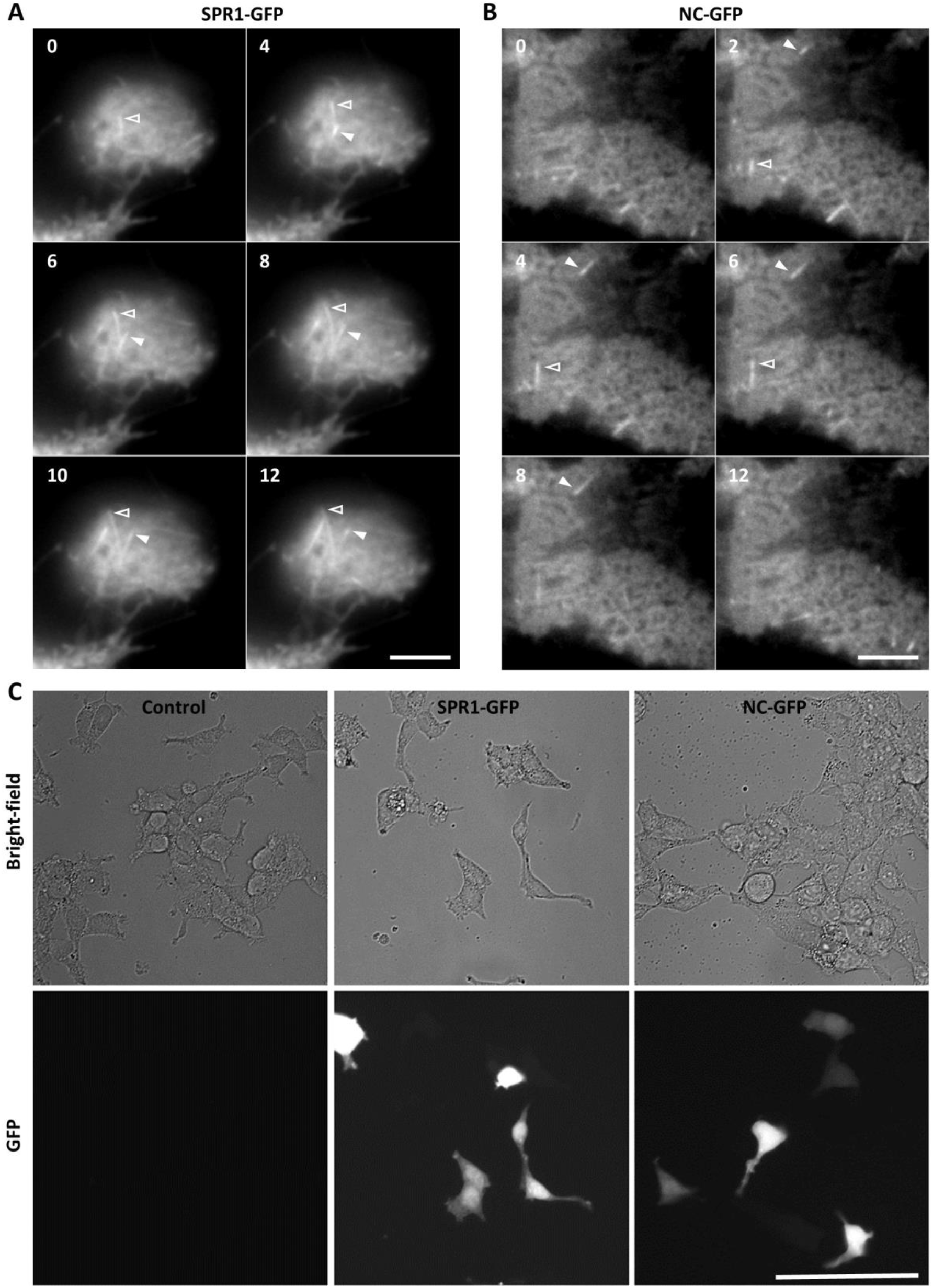
SPR1-GFP and NC-GFP label growing microtubule plus-end in heterologous systems. (A-B) Images of HEK-293 cells expressing SPR1-GFP (A) or NC-GFP (B). Arrowheads point to examples of labeled growing microtubule plus-ends. Numbers indicate time in seconds. Scale bar, 5um. (C) Bright-field (top row) and fluorescence (bottom row) images of HEK-293 cells either mock transfected (control) or transfected with SPR1-GFP or NC-GFP. Cells were imaged 48hr after transfection. Scale bar, 100um.

Live imaging of microtubule plus-ends in animal cells is commonly conducted by exogenously expressing fluorescent protein-tagged EB1 or EB3 (Ma et al., 2004; Mimori-Kiyosue et al., 2000; Salaycik et al., 2005; Stepanova et al., 2003; Yang et al., 2017). However, because EB proteins form the core of the microtubule plus-end complex (Lansbergen and Akhmanova, 2006), their ectopic expression can potentially disrupt the endogenous microtubule plus-end complex. As SPR1 is a plant-specific protein, it is likely to be more benign in animal cells. Since the NC construct is about half the size of SPR1 (~ 6 kDa) and minimally perturbs microtubule dynamics *in vitro*, we propose NC-GFP as a universal microtubule plus-end marker. Importantly, SPR1-GFP and NC-GFP did not visibly impair the viability and morphology of HEK-293 cells (Figure 4C), indicating that these proteins are not toxic to animal cells.

## MATERIALS AND METHODS

### Constructs

For recombinant protein expression, a commercially synthesized SPR1-GFP construct was cloned into pTEV plasmid between NcoI and XhoI restriction sites to obtain a C-terminal 6xHis tag. The N-GFP-C construct was assembled using recombinant PCR. For this, EGFP was PCR amplified with primers extending into the NSPR1 (5’-CAAAGCTCATTGGATTATCTCTTTGGTGGTGACGCTCCTATGGTGAGCAAGGGCGAG-3’) and CSPR1 regions (5’- TCCTTCAGCTCTGGCATAGTTGTTCTTGTACAGCTCGTCCATG-3’). Then, CSPR1 was PCR amplified with a forward primer containing 21bp of EGFP sequence (5’- GGCATGGACGAGCTGTACAAGAACAACTATGCCAGAGCTG-3’) and a reverse primer containing XhoI restriction site (5’-TATCTCGAGCTT GCCACCAGT GAAGAGAT AAT CCAAGGAT GATCCTCCTC-3’).

These PCR fragments were then mixed in 1:1 ratio and used as a template for a third PCR reaction with a forward primer consisting of the SPR1 N-terminal sequence and NcoI restriction site (5’-TATCCATGGGTCGTGGAAACAGCTGTGGTGGAGGTCAAAGCTCATTGGATTATC-3’) and a reverse primer containing XhoI restriction site (5’-TATCTCGAGCTTGCCACCAGTGAAGAGATAATCCAAGGATGATCCTCCTC-3’). The resulting PCR product was inserted into the NcoI and XhoI sites of the pTEV plasmid. The NC-GFP construct was obtained by fusing the N-and C-terminal domains of SPR1 in a two-step PCR. First PCR was using primers (5’-CAAAGCTCATTGGATTATCTCTTTGGTGGTGACGCTCCTAACAACTATGCCAGAGCTGAAG-3’ and 5—TATAGTCGACCTTGCCACCAGTGAAGAG-3’) and second PCR was using primers (5’-TATCCAT GGGTCGTGGAAACAGCT GTGGTGGAGGTCAAAGCTCATTGGATTATC-3’ and 5’-TATAGTCGACCTTGCCACCAGTGAAGAG-3’) and the first PCR product as template. The resulting PCR product was inserted upstream of GFP in the pTEV plasmid at NcoI and SalI site. GGG to AAA and PGGG to AAAA mutant versions of SPR1 were generated using site-directed mutagenesis. The SPR1 (Δ1-18)-GFP and SPR1 (Δ87-119)-GFP truncation constructs were generated by replacing full-length SPR1 with PCR amplified deletion fragments in the SPR1-GFP-6xHIS construct. All constructs created using PCR were verified by sequencing.

### Protein Purification

*E. Coli* Rosetta (DE3) cells were used for recombinant protein expression. Cells were grown to 0.6-0.8 OD600 at 30 °C and then induced using 0.15 mM IPTG at 16 °C for 16-20 hours. 6x-His-tagged proteins were isolated using Ni-NTA agarose beads. Purified proteins were desalted and exchanged into BRB80 buffer (80 mM piperazine-1,4-bis(2-ethanesulfonic acid), 1mM MgCl2, and 1mM EGTA, pH 6.8) supplemented with 10mM DTT and 50mM NaCl using PD-10 columns (GE healthcare). Protein concentration was measured using standard Bradford method. Proteins were aliquoted, snap frozen in liquid nitrogen and stored at −80 °C.

### Expression in Animal Cells

For expression in Human Embryonic Kidney (HEK)-293 cells, PCR amplified SPR1-GFP-6xHis, N-GFP-C-6xHis and NC-GFP-6xHis were cloned into the multiple cloning site of pLVX-puromycin plasmid. For HEK-293 cell transfection, 500ng of DNA and 2000ng polyethylenimine HCI MAX (Polysciences, Inc.) was added to 50ul of 300mM NaCl. This solution was mixed, incubated for 10 minutes at room temperature, and then added to the plated cells in culture media containing 10% fetal bovine serum in DMEM plus 1% Pen-Strep. Cells were imaged 24-48 hours after transfection.

### Total Internal Reflection Fluorescence Microscopy

For the *in vitro* reconstitution assays, flow chambers were prepared by attaching silanized coverslips to a glass slide using two layers of double-sided adhesive tape. GMPCPP (Jena Biosciences) stabilized microtubule seeds were prepared by polymerizing 50 μM porcine tubulin containing biotin-labeled (1:14.5) and rhodamine-labeled (1:14.5) porcine tubulins (Cytoskeleton, Inc.) in the presence of 1 mM GMPCPP at 37 °C for 30 min. The polymerized microtubules were then collected by centrifugation at 25,000g for 20 min, resuspended in 50 μL warm BRB80 containing 1 μM GMPCPP, and fragmented by passing through a 100 μL Hamilton syringe 4-5 times. Flow chambers were first coated with 20% anti-biotin antibody (Sigma Aldrich) followed by blocking with 5% Pluronic for 5 min each. Then 300 nM GMPCPP-stabilized seeds were flown in and allowed to bind to the anti-biotin antibody for 5 min. Microtubule polymerization was initiated by flowing in a mixture of 20 μM 1:25 rhodamine-labeled porcine tubulin, 1% methyl cellulose (4000 cP, Sigma-Aldrich), 50 mM DTT, 2 mM GTP, oxygen scavenging system (250 mg/ml glucose oxidase, 35 mg/ml catalase and 4.5 mg/ml glucose) and different concentrations of SPR1 or mutant proteins as indicated in the text. 5 mW 488-nm and 5 mW 561-nm diode-pumped, solid-state lasers were used to excite GFP and rhodamine, respectively. Images were collected by a 100X (NA 1.45) objective with 2x tube lens and a back-illuminated electron-multiplying CCD camera (ImageEM; Hamamatsu) at 2.5 s intervals and for a total of about 5 minutes.

For single-molecule imaging experiments, we used 100 nM, 150 nM and 250 nM of SPR1-GFP, NC-GFP and N-GFP-C proteins, respectively. Images were captured at 10 frames per second in streaming mode. A snapshot of microtubules was taken just before and immediately after the streaming mode acquisition to document microtubule growth.

### Data Analysis

Microtubule dynamics and single molecule kinetics were quantified from kymographs of individual microtubules generated using the Fiji ImageJ package (Schindelin et al., 2012). Microtubule growth and shortening rates were calculated for individual growth and shortening phases. Rescue frequency was calculated by dividing the sum of the number of transitions from shrinkage to growth by the time spent shrinking. Catastrophe frequency was calculated by dividing the sum of the number of transitions from growth to shrinkage by the time spent growing. To determine the microtubule transition frequency, a switch from growth to shortening or vice-versa was considered as a transition event. When a microtubule depolymerized all the way back to the GMPCPP seed followed a pause of 5 or more frames, subsequent microtubule growth was considered as an independent microtubule growth event and not a transition event of the previous microtubule. SPR1-GFP signal intensity plots were created using the plot profile tool in Fiji. Dot plots and statistical analyses were conducted using GraphPad Prism. All data points were included in the statistical analyses.

## ACKNOWLEDGEMENTS

We thank Marcus Mahar and Dianne Duncan for help with the animal cell experiments. We thank Rashmi Nanjundappa and Moe Mahjoub for providing the pLVX-puromycin plasmid. Funding was provided by grants from the National Science Foundation CAREER program (1453726) and National Institute of Health (R01 GM114678) to R.D.; and the National Institute of Health (R01 NS082446) to V.C.

## SUPPLEMENTAL INFORMATION

**Supplemental Table 1.** Effect of SPR1-GFP, N-GFP-C and NC-GFP on microtubule plus-end dynamics.

| | Growth Rate,<br>$\mu\text{m s}^{-1}$ (n) | Shortening Rate,<br>$\mu\text{m s}^{-1}$ (n) | Catastrophe Frequency,<br>events $\text{s}^{-1}$ (n) | Rescue Frequency,<br>events $\text{s}^{-1}$ (n) |
| --- | --- | --- | --- | --- |
| **[ SPR1 ]** |  |  |  |  |
| 0 nM | $0.052 \pm 0.013$<br>(67) | $0.489 \pm 0.172$<br>(31) | $0.005 \pm 0.009$<br>(52) | $0.031 \pm 0.034$<br>(27) |
| 50 nM | $0.046 \pm 0.009$<br>(147) | $0.501 \pm 0.213$<br>(68) | $0.004 \pm 0.005$<br>(98) | $0.061 \pm 0.072$<br>(58) |
| 200 nM | $0.043 \pm 0.008$<br>(229) | $0.576 \pm 0.414$<br>(134) | $0.004 \pm 0.003$<br>(139) | $0.054 \pm 0.071$<br>(106) |
| 500 nM | $0.032 \pm 0.006$<br>(276) | $0.495 \pm 0.252$<br>(177) | $0.009 \pm 0.014$<br>(160) | $0.082 \pm 0.105$<br>(129) |
| **[ N-GFP-C ]** |  |  |  |  |
| 0 nM | $0.035 \pm 0.006$<br>(156) | $0.478 \pm 0.256$<br>(90) | $0.005 \pm 0.007$<br>(106) | $0.056 \pm 0.096$<br>(67) |
| 50 nM | $0.040 \pm 0.041$<br>(218) | $0.525 \pm 0.288$<br>(122) | $0.007 \pm 0.021$<br>(144) | $0.064 \pm 0.085$<br>(98) |
| 200 nM | $0.039 \pm 0.064$<br>(199) | $0.528 \pm 0.296$<br>(121) | $0.007 \pm 0.013$<br>(129) | $0.066 \pm 0.118$<br>(101) |
| 500 nM | $0.035 \pm 0.005$<br>(188) | $0.518 \pm 0.478$<br>(111) | $0.005 \pm 0.006$<br>(117) | $0.065 \pm 0.068$<br>(90) |
| **[ NC-GFP ]** |  |  |  |  |
| 0 nM | $0.043 \pm 0.008$<br>(86) | $0.634 \pm 0.322$<br>(39) | $0.006 \pm 0.026$<br>(58) | $0.087 \pm 0.114$<br>(34) |
| 500 nM | $0.038 \pm 0.007$<br>(127) | $0.542 \pm 0.230$<br>(72) | $0.006 \pm 0.012$<br>(84) | $0.048 \pm 0.073$<br>(57) |
| 1000 nM | $0.042 \pm 0.006$<br>(123) | $0.547 \pm 0.183$<br>(62) | $0.005 \pm 0.015$<br>(77) | $0.058 \pm 0.066$<br>(49) |

Values are mean ± s.d. (n = number of events)

**Supplemental Movie 1**

SPR1-GFP localizes to growing microtubule plus-ends *in vitro.* Time is shown in seconds.

**Supplemental Movie 2**

N-GFP-C localizes to growing microtubule plus-ends *in vitro.* Time is shown in seconds.

**Supplemental Movie 3**

NC-GFP localizes to growing microtubule plus-ends *in vitro.* Time is shown in seconds.

**Supplemental Movie 4**

SPR1-GFP localizes to growing microtubule plus-ends in HEK-293 cells. Time is shown in seconds.

**Supplemental Movie 5**

NC-GFP localizes to growing microtubule plus-ends in HEK-293 cells. Time is shown in seconds.

